# Notch signaling pathway in tooth shape variations

**DOI:** 10.1101/2022.09.30.510272

**Authors:** Thimios A. Mitsiadis, Pierfrancesco Pagella, Helder Gomes Rodrigues, Alexander Tsouknidas, Liza L. Ramenzoni, Freddy Radtke, Albert Mehl, Laurent Viriot

## Abstract

Evolutionary changes in vertebrates are linked to genetic alterations that often affect tooth-crown shape, which is a criterion of speciation events. The Notch pathway is highly conserved between species and controls morphogenetic processes in most developing organs, including teeth. Epithelial loss of the Notch-ligand Jagged1 in developing mouse molars affects the location, size and interconnections of their cusps that lead to minor tooth-crown shape modifications convergent to those observed along Muridae evolution. These alterations are due to the modulation of numerous genes, where Notch signaling is a hub for essential morphogenetic networks. A three-dimensional metamorphosis approach allowed tooth morphology prediction in individuals carrying Jagged1 mutations. These results shed new light on Notch/Jagged1-mediated signaling as one of the crucial components for dental variations in evolution.

**Significance statement:** Dental microevolution changes in vertebrates are regulated by the Notch signaling pathway.

## Results and Discussion

Identifying the molecular mechanisms that have driven evolutionary changes in tissues and organs is a critical challenge in current biology. Teeth are the most mineralized tissues in vertebrates and therefore constitute the best-preserved part of the skeleton following fossilization. Consequently, studies of mammalian evolution often rely on the analyses of tooth shape, since subtle changes in tooth crown morphology usually constitute a criterion of speciation events. The remarkable diversity of tooth crown shapes results from differences in number, position, arrangement and interrelation of cusps, as well as on the dimensions of the dental crown. Variations in these traits reveal a wide range of adaptations that occurred in relation to numerous episodes of diversification over 200 million years of mammal evolutionary history (*1, 2*). Hence, the study of the genetic regulation of tooth morphology is important to understand the mechanisms underlying changes in tooth crown shape during evolution. In this context, the mouse dentition is one of the most widely used mammalian models in paleo-evo-devo investigations (*3-6*).

Mammals develop species-specific dentitions whose form and function are directly related to the activation of defined epithelial and mesenchymal signals at various locations of the developing tooth germs (*7*). A combination of key signaling molecules, including BMP (<u>B</u>one <u>M</u>orphogenic <u>P</u>rotein), FGF (<u>F</u>ibroblast <u>G</u>rowth <u>F</u>actor), Shh (<u>S</u>onic <u>h</u>edge<u>h</u>og) and Wnt families, are produced and secreted by the developing dental tissues. These molecules are also expressed during specific stages of odontogenesis in restricted areas of the dental epithelium that represent tooth exclusive signaling centers comparable to those localized in a variety of other developing organs in vertebrates (*7, 8*). These signals regulate the proliferative activity of epithelial and mesenchymal cells, leading to dental epithelial folding and formation of cusps, which constitute the earliest developmental sign of species-specific tooth patterning. Spatial arrangement and interconnection of cusps is closely linked to the diversity and evolution of dietary habits (*8*).

The Notch signaling pathway encompasses a group of evolutionary conserved trans-membrane protein receptors known to be involved in tooth formation and morphology (*9-11*). Four mammalian Notch homologues (Notch1, Notch2, Notch3 and Notch4), which interact with five trans-membrane-bound ligands (Jagged1, Jagged2, Delta1, Delta-like3 and Delta-like4), have been identified (*12, 13*). These molecules have been shown to play crucial roles in binary cell-fate decisions mediated by the lateral inductive cell signaling in many developmental systems. In humans, mutations in the *NOTCH1* and *NOTCH3* are associated with a neoplasia (T-cell acute lymphoblastic leukemia lymphoma) and CADASIL (cerebral autosomal dominant arteriopathy with subcortical infarcts and leukoencephalopathy), respectively (*14*). *JAGGED1* mutations have been associated with Alagille syndrome, an inherited autosomal dominant disorder that affects the face and various organs and tissues, including liver, heart, axial skeleton and eyes (*15, 16*). The functional role of Notch molecules has been investigated in detail by their targeted inactivation in mice. *Notch1*^*-/-*^(*17*), *Notch2*^*-/-*^ (*18*), *Jagged1*^*-/-*^ (*19*), *Jagged2*^*-/-*^ (*20*), *Delta1*^*-/-* 20,^(*21*) homozygous null mice die either at early embryonic stages or after birth. Notch deregulation in mice severely affects the development of the kidney, heart, intestine, eyes, somites, as well as neurogenesis and angiogenesis (*12, 17-21*). Components of the Notch pathway are also expressed during odontogenesis and our previous findings have shown that *Jagged2*-mediated Notch signaling is required for proper tooth morphology (*20*).

Our previous studies have shown that *Jagged1* is expressed from the very first stages of odontogenesis in the developing dental epithelium (*9*). Loss of *Jagged1* is embryonically lethal due to defects in the embryonic and yolk sac vasculature remodeling (*19*). Therefore, to investigate the role of this gene in dental epithelium we used a tissue-specific deletion strategy (*K14Cre;Jagged1*^*fl/fl*^; GenBank accession number AF171092). We first examined the effects of *Jagged1* loss in tooth epithelium and its involvement in tooth crown shape modifications. Teeth of *K14Cre;Jagged1*^*fl/fl*^ mice exhibited crown morphological differences when compared to teeth of wild-type (WT) mice, mainly regarding the location and interconnection of the cusps (Fig. 1Aa; red arrows). The most striking variations on the upper molars (i.e., M^1^, M^2^, M^3^) were observed in their first cusp (c^1^ cusp). In WT mice, the c^1^-c^2^ connection of the M^1^ seen in side-view was usually achieved by a high V-shaped crest, which contrasts to the frequently observed (in approximately 60 % of analyzed samples) low U-shaped profile in M^1^ of the *K14Cre;Jagged1*^*fl/fl*^ mice (Fig. 1Aa, Ab; red lines). This implies that the c^1^-c^2^ connection was weak to absent in the *K14Cre;Jagged1*^*fl/fl*^ mutant mice (Tab. 1). This variation was confirmed by height measurements of the c^1^-c^2^ connection (Fig. 1B; d1), which were significantly different between the two cohorts. In addition, these measurements revealed that the spacing between c^1^ and c^2^ cusps was significantly greater in the M^1^ of mutant mice (Fig. 1B; d2). The c^1^ cusp of the M^1^ had a more linguo-distal position in the majority of the *K14Cre;Jagged1*^*fl/fl*^ mice, and this feature explained its greater spacing from c^2^. On the M^2^ of mutant mice, the vestibular extension of the cusp c^1^ spur was absent in about 60% of specimens (Fig. 1Aa; Tab. 1). The c^1^ cusp of the M^3^ was sometimes reduced and tended to merge with the c^4^ cusp in 30% of the *K14Cre;Jagged1*^*fl/fl*^ specimens (Fig. 1Aa; Tab. 1). The central cusps (c^5^ and c^8^) of the mutant M^1^ and M^2^ had a rather angular or even pointed mesial edge, while this part was always smooth in the molars of WT mice (Fig. 1Aa). This latter morphotype occurred in approximately 60% of the upper molars of *K14Cre;Jagged1*^*fl/fl*^ mice (Tab. 1), but was not always simultaneously present on each cusp. The lower molars (i.e. M_1_, M_2_, M_3_) of the mutant mice showed little variation compared to the upper molars, yet the c_1_ and c_2_ cusps of the mutant M_1_ appeared closer to each other compared to the WT M_1_ (Fig. 1Aa). Three measurements revealed significant differences in d3-d5 mean lengths (Tab. 2) and confirmed that c_1_ and c_2_ cusps tended to partly merge in M_1_ of mutant mice, while the cusp c_1_ was less protruding.

**Figure 1.**
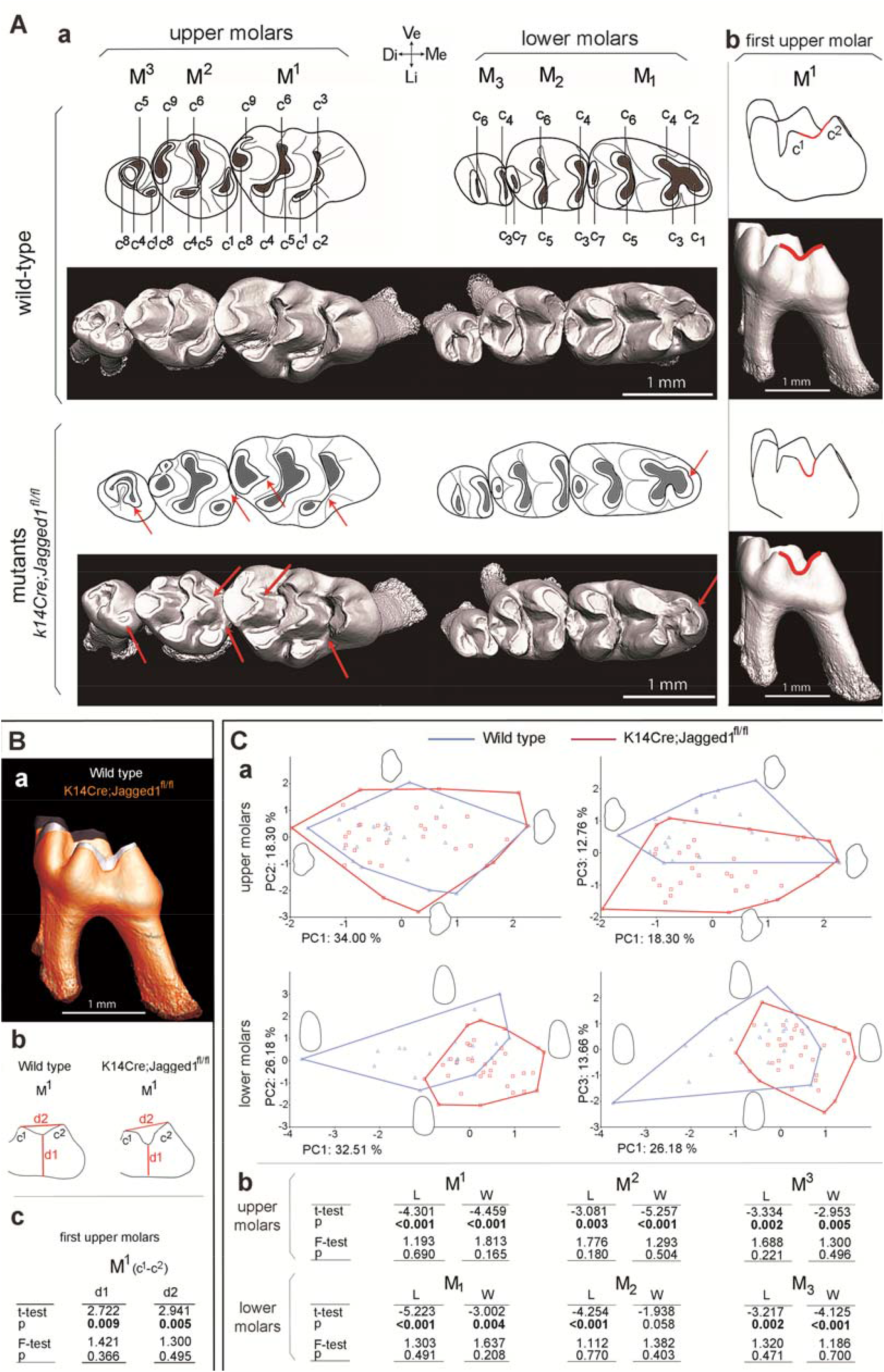
Epithelial loss of Jagged1 affects the molar dental morphology. (A) MicroCT analysis showing the morphology of upper and lower molars of wild-type (WT) and *K14Cre;Jagged1fl/fl* mutant mice (both occlusal and side views). (Aa) Morphological variations in location, size and connection of the cusps in molars of the *K14Cre;Jagged1fl/fl* mice are indicated by red arrows. (Ab) WT first upper molar (M^1^) showing a V-shape c^1^-c^2^ connection profile, and mutant M^1^ showing a U-shape profile (red lines). In mutant mice, the lower first molars (M_1_) show in the mesial part mild fusion between the c_1_ and c_2_ cusps. (B) Comparison of the M^1^ crowns in WT and Jagged2 mutants. (Ba) MicroCT analysis showing smaller crowns in mutant M^1^ (orange pseudo-color) when compared to the crowns of WT M^1^ (grey color). (Bb,c) Comparison of the distance between the intercusp groove and the basis of the crown (d1) and the distance between c^1^ and c^2^ (d2) in M^1^ of WT and mutants. (C) Principal component analyses comparing the variation of dental outline of M^1^ and M_1_ between WT (blue color) and *K14Cre;Jagged1fl/fl* (red color) mice. Abbreviations: M, Mesial part; D, Distal part; V, Vestibular part; L, Lingual part.

**Table 1.**
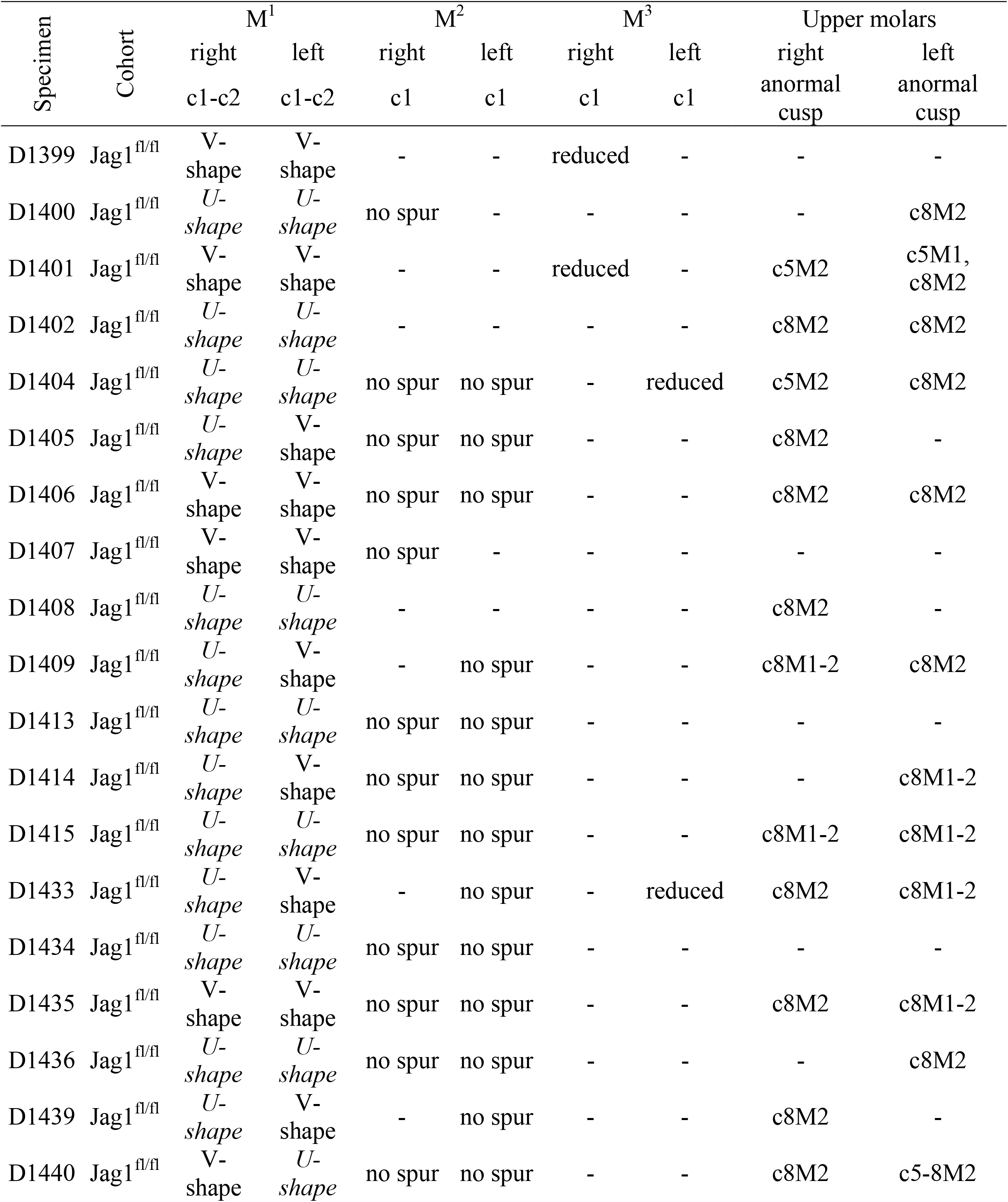
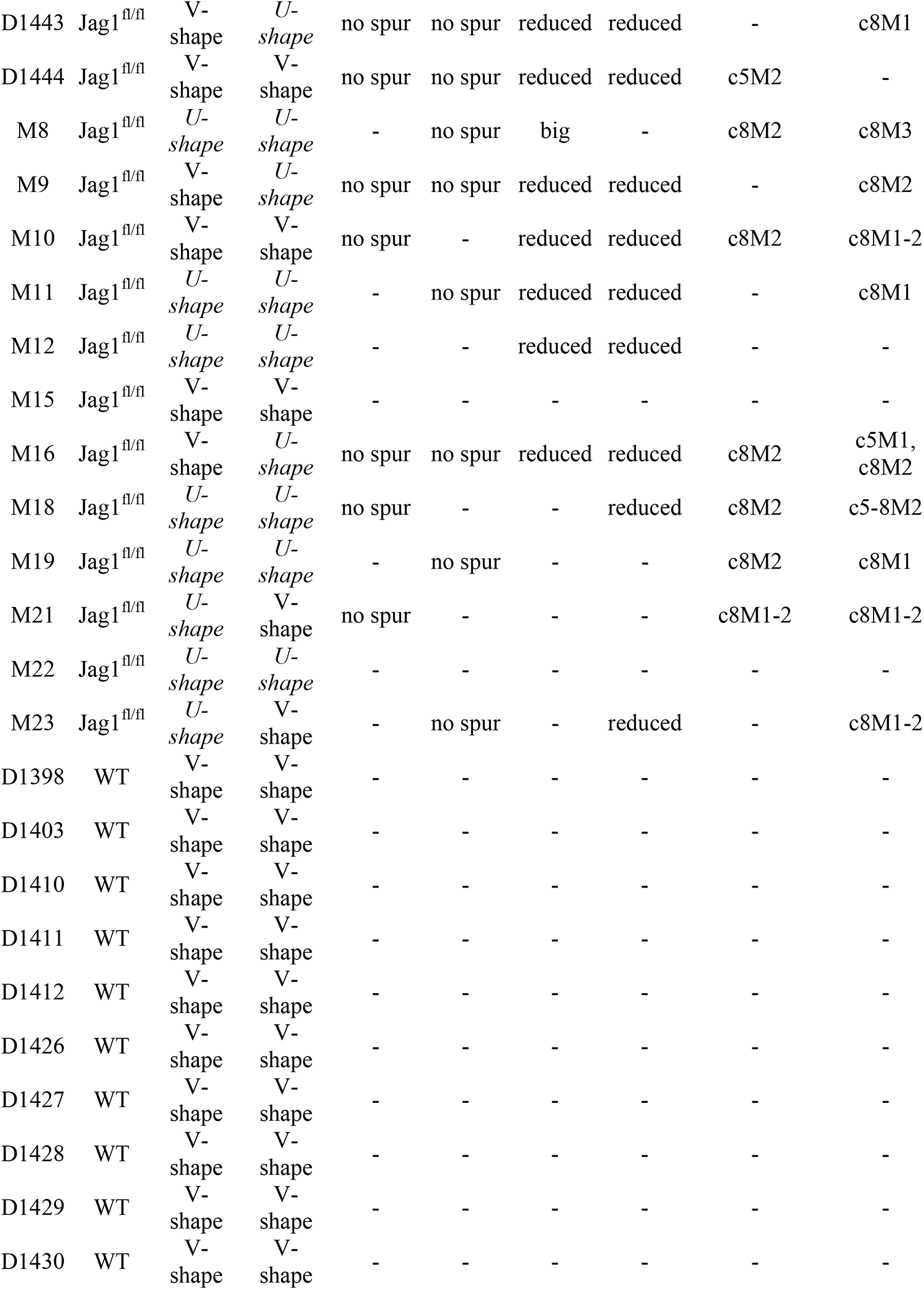
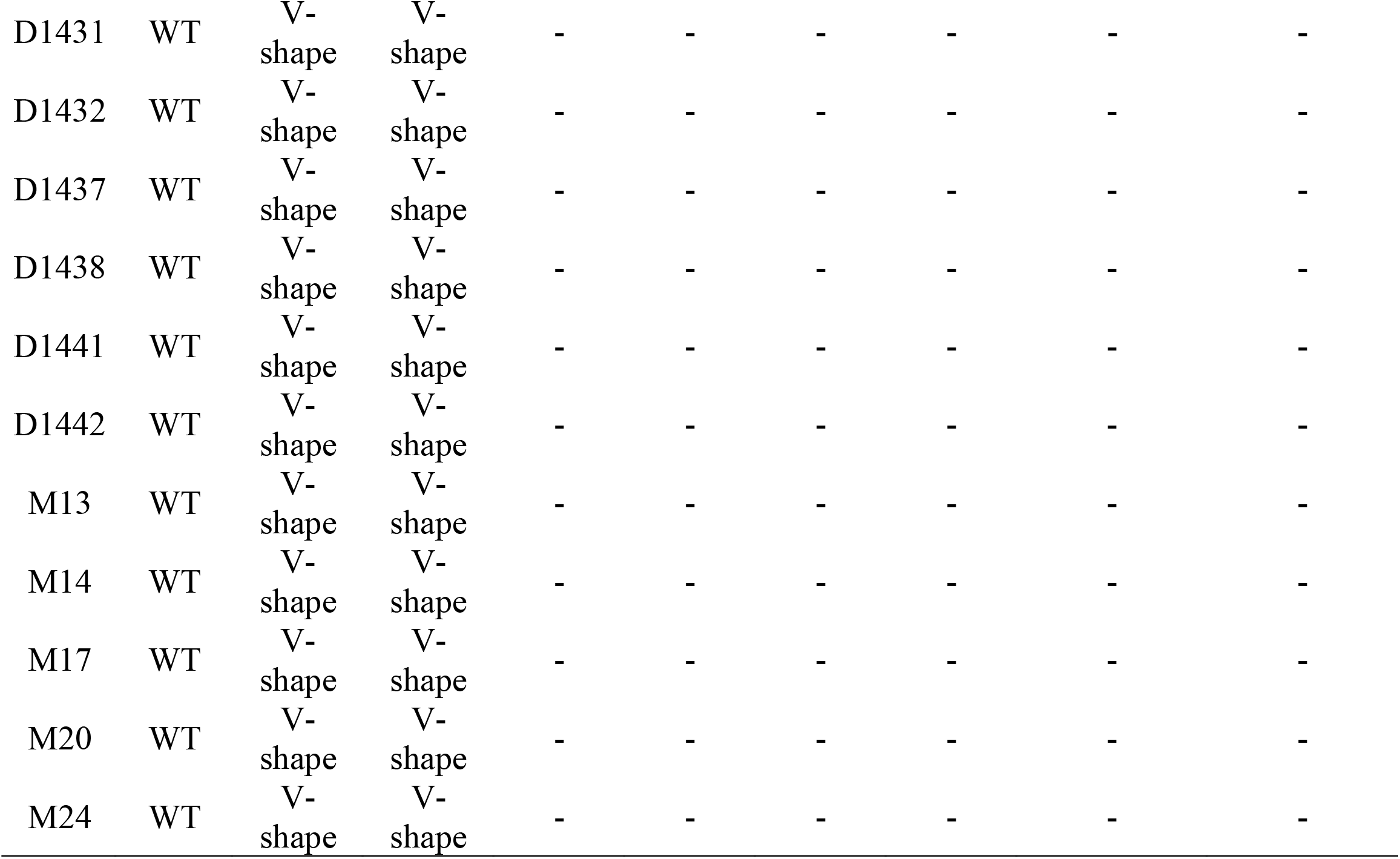

**Table 2.**
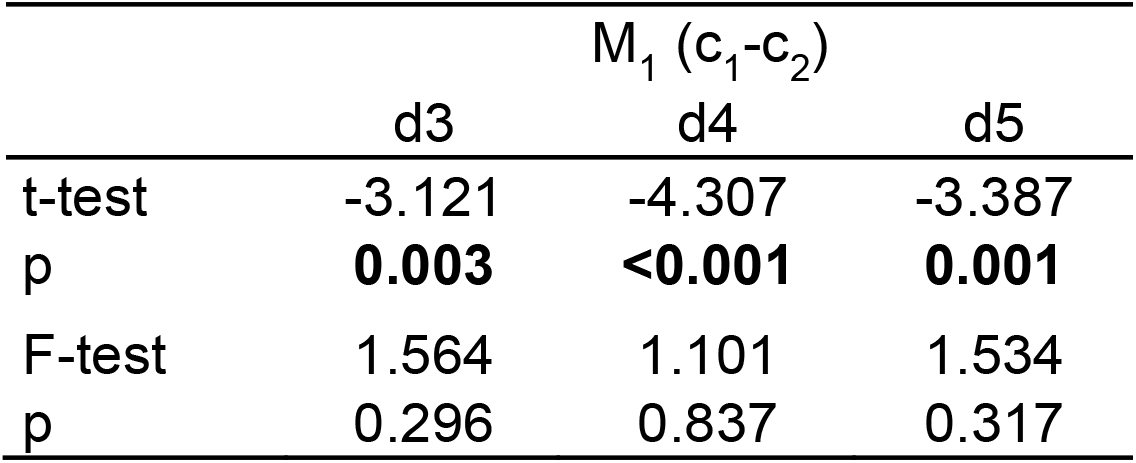

The first two axes of the PCA on M^1^ outlines were poorly discriminant because WT and *K14Cre;Jagged1*^*fl/fl*^ specimens plotted together, but the third axis was more discriminant although morphospaces partly overlapped (Fig. 1Ca). The main dental trend expressed on the third component was the variation of the cusp c^1^ location, according to extreme outlines on the negative and positive sides. This cusp was indeed located in a more distal-lingual position in the extreme outline of the negative side where a majority of *K14Cre;Jagged1*^*fl/fl*^ specimens plots. The results of the principal component analysis (PCA) were also confirmed by a multivariate analysis of variations (MANOVA), pointing out a significant difference in M^1^ outline between WT and *K14Cre;Jagged1*^*fl/fl*^ mice (Tab. 3). Contrary to M^1^, the first component of the PCA on M_1_ outlines represented the most discriminant axis, and the second one was more discriminant than the third (Fig. 1Ca). Nonetheless, the M_1_ morphospaces overlapped as well. The main difference between WT and *K14Cre;Jagged1*^*fl/fl*^ was linked to the mesial protrusion of the c_1_ cusps, which was less important on extreme outlines from the negative to the positive side of the first component. A significant difference between the two cohorts was also displayed by the MANOVA (Tab. 3). Mean sizes (L and W) of all molars were significantly lower in *K14Cre;Jagged1*^*fl/fl*^ mice compared to WT specimens (Fig. 1Cb; t-test), while there was no significant difference concerning the variances (Fig. 1Cb; Fischer’s F-test).

**Table 3.**
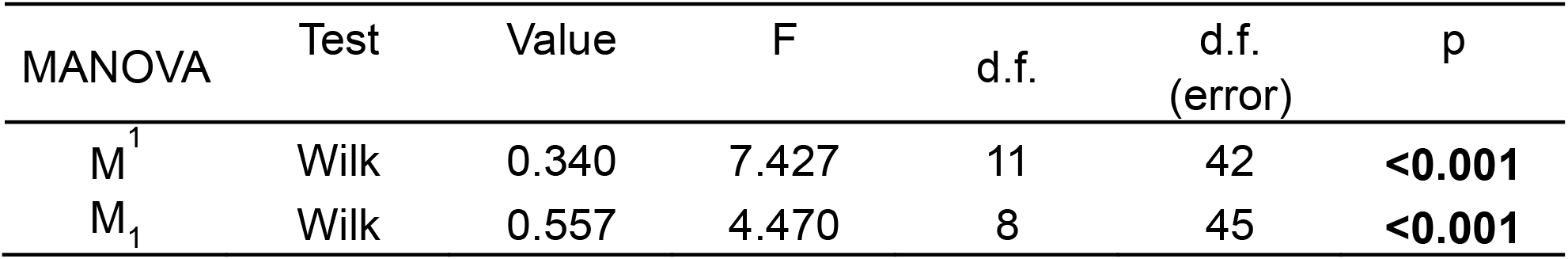

The characteristics of the *K14Cre;Jagged1*^*fl/fl*^ mouse molars provide relevant information in terms of tooth crown microevolution in rodents. The evolution of the rodent dentition is well understood from an abundant and intensively studied fossil record (*6, 22*). The transition from ancestral to descendant species within most evolutionary lineages usually involves minor modifications of tooth shapes. The subtle changes in the interconnections between cusps (Fig. 1A), their varying frequency, and the overlapping tooth size range are reminiscent of cases of dental variations among muroid rodents. Pleistocene and Holocene species of field mice (*Apodemus*) have been shown to possess M^1^ with variable interconnections between the c^1^ and c^2^ cusps, as well as variable interrelations between the c_1_ and c_2_ cusps on the M_1_, and these variations occur both at intraspecific and close interspecific levels (*23*). Variations in the interrelationships between mesial cusps of M^1^ and M_1_ have also been described in different Miocene species of *Democricetodon* (*24*) and *Megacricetodon* (*25*). In most of these examples, the tooth size range remains nearly identical.

The loss of *Jagged1* function in dental epithelium can thus cause morphological changes in the same cusps that are modified during murid evolution. Consequently, an alteration in *Jagged1* expression might be responsible for subtle tooth crown modifications that underlie processes of dental phenotype adaptation during population splitting or even speciation events over evolution. Tooth morphogenesis requires the orchestrated activity of several molecular networks, and the Notch pathway plays a pivotal role in this process, acting as a hub regulator of main signaling pathways (*26*). To understand how the epithelial deletion of *Jagged1* modulates the whole dental developmental program we compared the transcriptome of first molars isolated from E18 *K14Cre;Jagged1*^*fl/fl*^ embryos and WT littermates (Fig. 2A).

**Figure 2.**
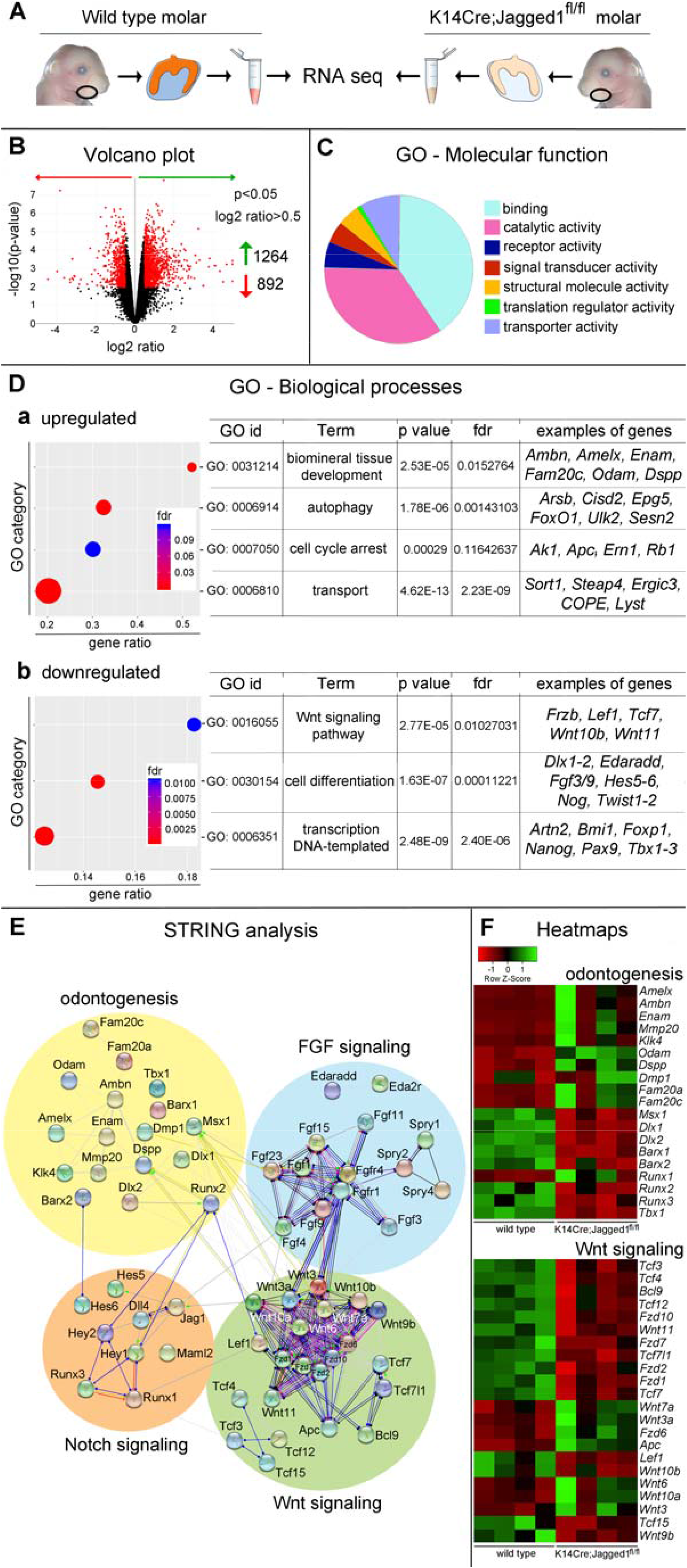
RNA sequencing analysis of first molars isolated from E18.5 WT and *K14Cre;Jagged1*^*fl/fl*^ mouse embryos. (A) Experimental approach. Molars were isolated from E18.5 *K14Cre;Jagged1*^*fl/fl*^ embryos and WT littermates. Their transcriptome was isolated via mRNA sequencing. (B) Volcano plot showing downregulated (red arrow) and upregulated (green arrow) genes in *K14Cre;Jagged1*^*fl/fl*^ mutant mice compared to the WT littermates. (C) Pie chart representation of unbiased Gene Ontology Analysis – Molecular Function. (D) Hypergeometric representation of unbiased GO – Biological Processes analysis showing the upregulated (Da) and downregulated (Db) genes in the *K14Cre;Jagged1*^*fl/fl*^ molars. (E) STRING analysis of selected protein networks whose gene expression was found to be significantly affected by *Jagged1* deletion in the dental epithelium. (F) Heatmaps showing the main deregulated genes involved in odontogenesis and in the Wnt signaling pathway in *K14Cre;Jagged1*^*fl/fl*^ molars.

RNA sequencing analysis showed a significant change in the expression of more than 2000 genes upon epithelial deletion of *Jagged1* (Fig. 2A, B). These genes encode diverse categories of proteins, including proteins linked to binding, catalytic and transported activities (Fig. 2C). Unbiased Gene Ontology (GO) Enrichment Analysis identified several networks affected by the loss of Jagged1 (Fig. 2D). We found a significant upregulation of genes involved in mineralization, autophagy, ion transport and cell cycle arrest (Fig. 2D, Fig. S3-8), which are critical processes for enamel and dentin formation (*27, 28*). Several genes encoding for proteins necessary for enamel formation, like *Amelx, Ambn, Enam, Mmp20*, and *Klk4* were upregulated in *K14Cre;Jagged1*^*fl/*fl^ molars (Fig. 2E). However, these results varied between the *K14Cre;Jagged1*^*fl/*fl^ molars (Fig. 2F). In addition, the expression of mesenchymal genes was also affected in *K14Cre;Jagged1*^*fl/fl*^ molars. For example, *Pax9, Barx1, Dlx1* and *Dlx2* were significantly downregulated, while genes associated with odontoblast differentiation such as *Dspp* and *Dmp1* were upregulated in the mutant molars (Fig. 2D, F). Collectively, these findings suggest a premature cytodifferentiation process in both dental epithelium and mesenchyme of *K14Cre;Jagged1*^*fl/fl*^ mice.

Several signaling pathways that are important for odontogenesis were also affected upon Jagged1 deletion, including the Wnt signaling pathway. We observed major downregulation of genes encoding Wnt11, Wnt10b, Wnt9b ligands, Frizzled receptors and Tcf transcription factors (Fig. 2C, D, E), while the genes encoding Wnt3a, Wnt6, Wnt7a and Wnt10a ligands, as well as Apc were upregulated (Fig. 2E). The perturbed expression levels of Wnt signaling molecules are expected due to the well-known genetic interaction with the Notch signaling pathway (*29-31*). The Wnt signaling is a powerful morphogen (*32*) and regulator of planar cell polarity (*33*), fundamental for tooth development and mutations in this pathway are associated with dental anomalies (*34-39*). Transcriptional changes were also observed in the FGF signaling pathway (Fig. 2E, F), namely downregulation of the pathway inhibitors *Spry1, Spry2* and *Spry4* (Fig. 2E, F; Fig. S1), mutations of which affect tooth number (*40*) and enamel formation (*41*).

Notch pathway members, such as *Jagged1, Hes5, Hes6, Lfng*, and *Maml2* were downregulated, while others, such as *Hey1* and *Dll4* were upregulated (Fig. 2D, E; Fig. S1). Despite the known correlation between Notch signaling and cell proliferation (*42*), no significant alterations were observed in specific networks linked to cell proliferation events. The upregulation of several cell cycle arrest-related genes further supports premature cytodifferentiation events in the mutant molars (Fig. 2Da).

To validate these results and analyze whether the site of expression of different genes and proteins was affected by *Jagged1* epithelial deletion, we performed real time-PCR analysis, *in situ* hybridization, and immunohistochemistry on cryosections of E18 *K14Cre;Jagged1*^*fl/fl*^ mouse embryos (Fig. 3). Real time PCR (RT-PCR) analysis confirmed *Jag1* downregulation as well as *Notch1, Notch2*, and *Hes5* downregulation in *K14Cre;Jagged1*^*fl/fl*^ molars (Fig. 3A). We further demonstrated overexpression of ameloblast (e.g. *Amelx* and *Ambn*) and odontoblast (e.g. *Dspp* and *Dmp1*) differentiation markers in *K14Cre;Jagged1*^*fl/fl*^ molars (Fig. 3A). RT-PCR also confirmed alterations in the expression of genes coding for components of the Wnt signaling pathway, namely the upregulation of *Wnt10a* and *Apc*, and downregulation of *Lef1, Tcf3* and *Fzd2* (Fig. 3A). *In situ* hybridization analysis in E18 WT molars showed intense *Jagged1* expression in cells of the inner dental epithelium, which was drastically reduced in this cell layer in E18 mutant molars (Fig. 3B). Although strong *Notch1* expression was observed in all cells of the stratum intermedium in WT molars, its expression was downregulated in cells located at the cusp territories of *K14Cre;Jagged1*^*fl/fl*^ molars (Fig. 3B). Similarly, *Notch2* expression was downregulated in cells of the outer enamel epithelium and stellate reticulum of E18 mutant molars, when compared to the gene expression in WT molars (Fig. 3B). The expression of *Jagged2* in inner dental epithelial cells was comparable between the mutant and WT molars (Fig. 3B), which suggests a *Jagged2* compensation for the loss of *Jagged1*, and can thus explain the mild morphological changes observed in the mutant molars. Immunohistochemistry confirmed the deregulation of the Notch signaling pathway in the dental epithelium of WT molars. Labelling against the active form of Notch1 (i.e. Notch1 intracellular domain) showed its distribution in cells of the stratum intermedium of E18 WT molars, contrasting the lack of immunostaining in mutant molars (Fig. 3C). Similarly, although a strong Hes1 staining was observed in the stratum intermedium of WT molars, in the cusps of the mutant molars the staining was not obvious (Fig. 3C). Hes5 staining was mainly detected in inner dental epithelium (preameloblasts), stellate reticulum and odontoblasts of the E18 WT molars (Fig. 3C). Comparison of Hes5 immunoreactivities between WT and mutant molars indicated a slight reduction of the staining in the inner dental epithelium and odontoblasts in the cusp areas (Fig. 3C). Amelogenin was distributed in preameloblasts located at the cusps of the WT molars, as well as in parts of the dental pulp (Fig. 3C). A similar pattern for Amelogenin was observed in the *K14Cre;Jagged1*^*fl/fl*^ molars, although the staining appeared more expanded when compared to that of WT teeth (Fig. 3C). Staining with the proliferation marker Ki67 showed mitotic activity in few cells of the stratum intermedium located in the cusps of the WT molars (Fig. 3C). In mutant teeth, increased Ki67 immunoreactivity was observed in cells of the stratum intermedium and some preameloblasts (Fig. 3C).

**Figure 3.**
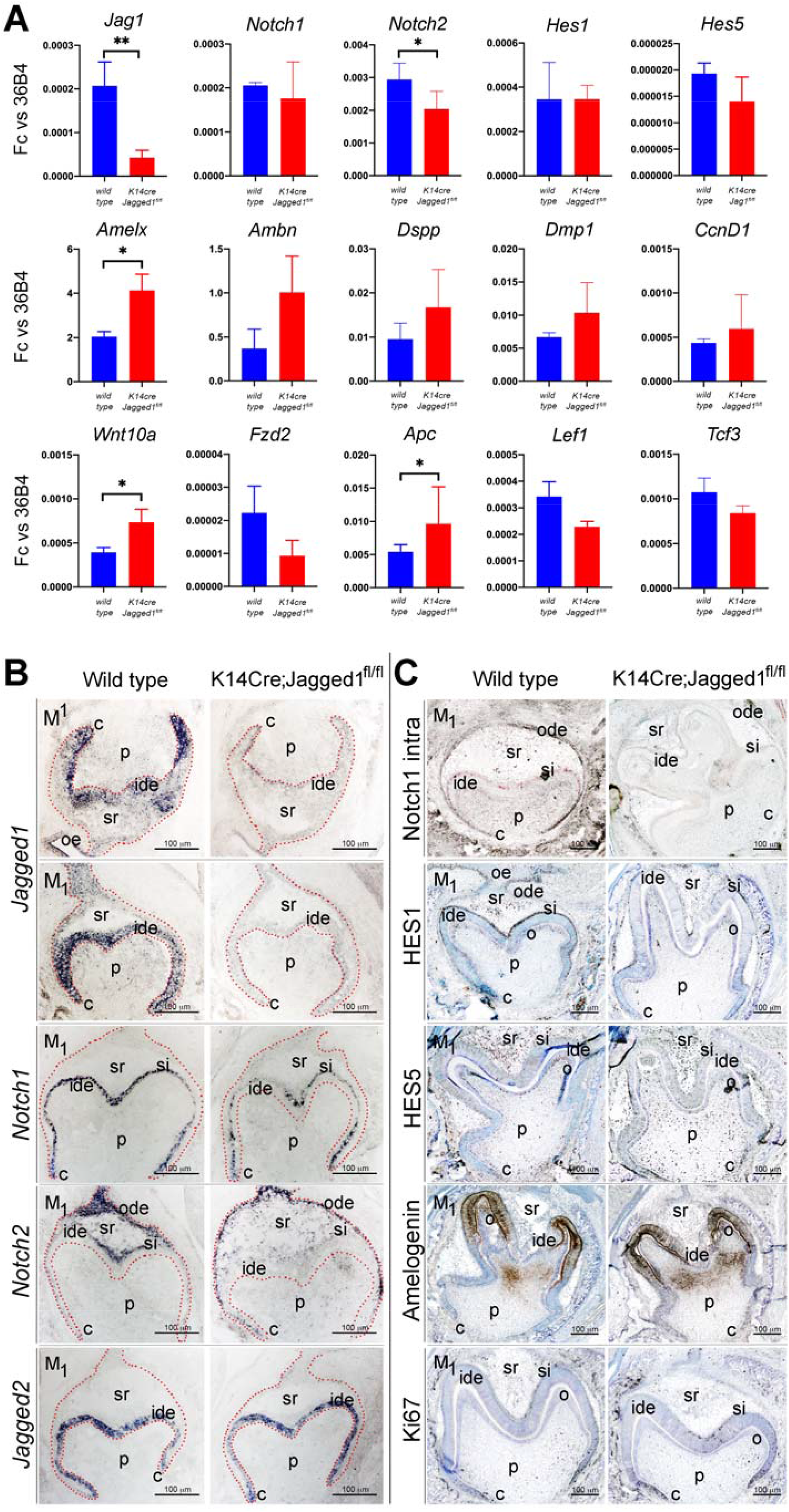
Expression analysis of genes and proteins in E18.5 WT and *K14Cre;Jagged1*^*fl/fl*^ teeth. (A) RT-PCR analysis of the expression of genes coding for components of the Notch pathway (i.e. *Jag1, Notch1, Notch2, Hes1, Hes5*), Wnt signaling pathway (i.e. *Wnt10A, Fzd2, Apc, Lef1, Tcf3*), various ameloblast (i.e. *Amelx, Ambn*) and odontoblast (i.e. *Dspp, Dmp1*) differentiation markers, and a proliferation marker (i.e. *CcnD1*), in both WT (blue color) and mutant (red color) first molars. (B) *In situ* hybridization showing alterations in the expression pattern of *Jagged1, Notch1, Notch2* and *Jagged2* in mutant first molars when compared to WT ones. (C) Immunohistochemistry showing modifications in the distribution pattern of the active form of Notch1 (Notch1 intra), Hes1 and Hes5 proteins, the enamel-specific Amelogenin protein and the nuclear Ki67 protein (cell proliferation marker) in *K14Cre;Jagged1*^*fl/fl*^ teeth when compared to WT teeth. Scale bars 100µm.

Although the extent to which genetic changes contribute to morphological variations in human dentition remains a fundamental question in evolutionary biology, the use of three-dimensional (3D) modeling techniques can provide some insight into potential diversities. Despite the obvious differences between mouse and human molars (in cusp number, size and interconnections as well as in relative dental proportions), different studies indicated some possible equivalences (*43*). Thus, the understanding of the molecular bases underlying fine variations in mouse dental morphology can help us to unravel the common factors that shaped teeth throughout evolution. On this basis, we investigated how the deregulation of Jagged1 function would affect the morphology of human molars. Based on mammalian evolution of common ancestral origin (*44*) and by using 3D metamorphosis techniques (*45*), the statistically significant variations between WT and mutant rodent teeth can be mathematically applied to the human teeth, thus allowing to predict the crown morphology in humans carrying Jagged1 mutations. For building this model, we opted for M^1^ where the most accentuated modifications were observed in the mutant mice. A polygonal surface technique (3D direct morphing, using ANSA by BETA CAE Systems S.A.) was applied to compute a translation matrix. The transformation vectors of the translation matrix contain all the important information that would allow approximation of the WT M^1^ crown morphology to the *K14Cre;Jagged1*^*fl/fl*^ one (Fig. 4Aa, b), while providing an overview of the required node displacements (Fig. 4Ac). These node displacements can be described by transformation vectors (Fig. 4Ad), which can be transferred to the human M^1^. Consequently, we superimposed WT mouse M^1^ and human M^1^ morphologies (Fig. 4B). Despite the fact that significant variations in mouse M^1^ also exist in its mesial part that is not comparable to the human M^1^, an analogue was drawn to the remaining variations in the distal part (containing the c^5^, c^6^, c^8^ and c^9^) of the mouse M^1^. The transformation matrix of each of these four cusps in the mouse M^1^, was applied to the 3D data set of the average human M^1^ (Fig. 4C), to predict tooth morphology in individuals carrying Jagged1 mutations (Fig. 4Cb, Da-d). The most significant morphological alterations were observed in the occlusal surface of M^1^, with a mesial-lingual/palatal shift of the hypocone and a mesial displacement of the metacone (arrows in Fig. 4Cb, 4Da, b). The morphologies of the protocone and paracone were only marginally affected by the mutation. The distal ridge of the mutant M^1^ is predicted to have a notable deeper and narrower profile (Fig. 4Dd) when compared to a typical M^1^ morphology (Fig. 4Dc). The distal-lingual/palatal groove was also computed as slightly widened in its mesial part (Fig. 4Db), while no significant changes were observed in the vestibular surface of the mutant M^1^.

**Figure 4.**
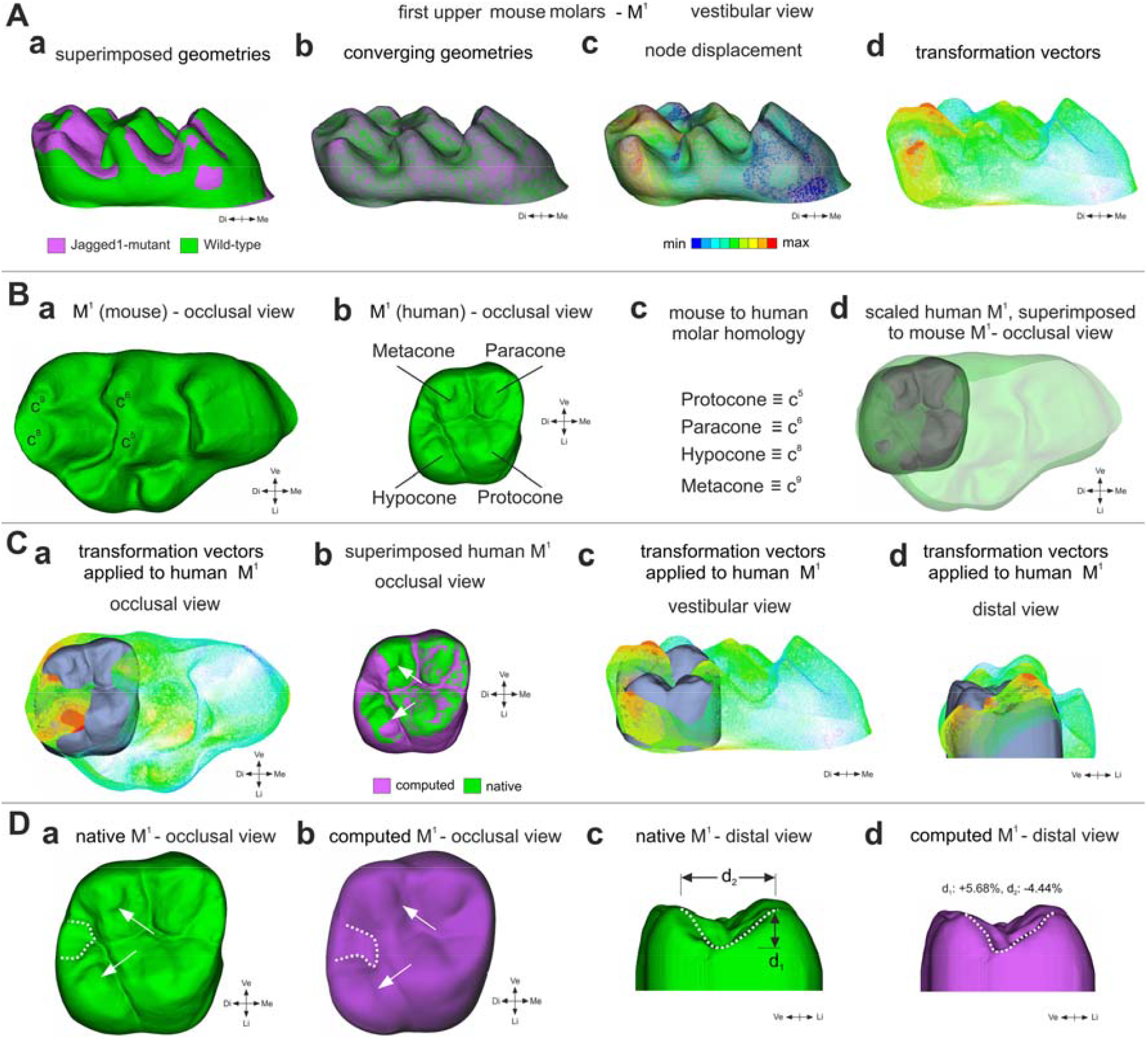
Prediction of tooth crown morphologies in humans carrying Jagged1 mutations. (Α) Compared morphologies of WT and *K14Cre;Jagged1*^*fl/fl*^ M^1^. (Aa) Pre-processing (scaling, positioning and alignment) of proxy geometries, illustrating the affinity of source and target surfaces of rodent M^1^. Green color represents WT teeth, violet color indicated mutant teeth. (Ab) Conversion of the morphed WT M^1^, approximating the *K14cre;Jagged1*^*fl/fl*^ M^1^ geometry. (Ac) Magnitude of nodes movement required for the approximation of the WT M^1^ geometry to the *K14cre;Jagged1*^*fl/fl*^ M^1^ geometry. (Ad) Indicative view of the transformation vectors from WT M^1^ to *K14cre;Jagged1*^*fl/fl*^ M^1^. (B) Tooth homology between mouse and human M^1^ molars. (Ba) Occlusal view of mouse WT M^1^ indicating the comparable to the human M^1^ cusps. (Bb) Occlusal view of human M^1^ indicating the four cusps. (Bc) Nomenclature homologation in-between the human M^1^ and mouse M^1^ cusps. (Dd) Scaled human M^1^ superimposed on WT mouse M^1^. (C) Transformation vectors of the morphing process representing the magnitude of the source nodes’ (WT M^1^) movement during their approximation of the *K14Cre;Jagged1*^*fl/fl*^ M^1^ geometry. (Ca, c, d) Transformation vectors superimposed on human M^1^. (Cb) Superimposed native (green color) and computed (violet color) human M^1^. (D) Comparison of the native human M^1^ and predicted human mutant M^1^. (Da, b) Native (green color in Da) and computed (violet color in Db) human M^1^ indicating statistically significant changes in cusps (arrows) and the inter-cusp space (dotted lines). Occlusal view. (Dc, d) Native (green color in Dc) and computed (violet color in Dd) human M^1^ showing a notable deeper and narrower profile of the distal ridge in mutant teeth (dotted lines in Dd) compared to the native teeth (dotted lines in Dc). Distal view.

Taken together, our results suggest that the systematic effect of *Jagged1* deletion provides a basis for dental variations in evolution. At a transcriptome level, *Jagged1* epithelial loss induces important modifications of numerous genes that are involved in an important and diverse number of signaling pathways. Loss of Jagged1 in mice did not result in dramatic changes in the activity of these signaling networks, but rather fine and discrete modulations, which still significantly impacted crown morphology. This indicates that Jagged1 fine tunes a sensitive balance between various gene networks whose interactions govern organogenesis. Disturbance of this genetic equilibrium might ultimately lead to the generation of subtle morphological modifications in most tissues and organs. Wnt signaling is involved in the establishment of planar cell polarity (PCP) that plays a crucial role in tissue patterning during all developmental stages (*33*). Significant alterations in the Wnt signaling pathway in the dental epithelium of Jagged1 mutants suggest that tooth-crown morphological changes are mostly due to PCP dysfunction. This hypothesis is further reinforced by the deregulation of genes that are crucial for the establishment of PCP, such as *Vangl2* and *Celsr1* (*33*), as well as by the altered expression patterns of *Notch1* and *Notch2* in mutant teeth. Misexpression of the Notch receptors and ligands in dental epithelium, in combination with the deregulation of various Notch downstream effectors, will also affect the fate of dental epithelial cells (*12*). The upregulation of genes involved in ameloblast (e.g. *Amelx, Ambn* and *Enam*) and odontoblast (e.g. *Dspp* and *Dmp1*) differentiation, together with the upregulation of genes associated with cell-cycle arrest, indicates premature cytodifferentiation in mutant molars. As a consequence, the accelerated deposition of enamel and dentin by the ameloblasts and odontoblasts, respectively, will prematurely stabilizes tooth-crown shape in the mutants. Premature cytodifferentiation events in mutant molars could thus lead to the observed reduced size and subtle morphological alterations of their crown. Such dental morphology does not resemble any particular murid rodent or any specific dental pattern that can be found in the evolution of Muridae. This is different from changes observed upon *Fgf3* deregulation in mice that greatly affected tooth morphology (*4*). Therefore, the present results should be evaluated from a microevolutionary perspective rather than from a macroevolutionary viewpoint. Indeed, the trends observed here for some characters (e.g. c^1^-c^2^ connection, c^1^-c^2^ complex), their variable frequency, and the overlapping of both size range and shape reminds of cases of wild sibling species of Muroidea (*46, 47*). It was shown that two extant field mice (i.e. *Apodemus sylvaticus* and *Apodemus flavicollis*) share dental morphologies and nearly the same size ranges (*46*). Moreover, the extinct “hamster-like” species, *Eucricetodon asiaticus* and *Eucricetodon jilantaiensis*, from the Oligocene of Ulantatal (Inner Mongolia, China), displayed some overall shape variations even closer to the *K14Cre;Jagged1*^*fl/fl*^ and WT case. The size range of these species is indeed nearly identical, they share intermediary dental morphotypes, their global shapes overlap and their main size distinction relies on mean L/W ratio (*47*). Consequently, modulation of Jagged1-mediated signaling could be the driving force for minor dental modifications that contribute to speciation phenomena, despite the reported negative effect of a haploinsufficiency of this gene in some organs (*16, 48-50*). Mutations morphologically similar to Jagged1 deletion shed a completely new light on the mechanisms of development, which could reconcile microevolutionary and macroevolutionary processes.

## Acknowledgments

We thank the staff of the Institute of Oral Biology and Dr M. Alexiou (University of Zurich) for technical assistance with the mice. The authors would like to thank BETA CAE Systems A.G. for providing them with the CAE pre-processor ANSA, used during surface modeling and morphing of the teeth. The authors would also like to thank Dr Giancarlo Russo from the Functional Genomics Center Zurich (University of Zurich) for helping with the bioinformatics analysis of the RNA sequencing data, and Dr Anamaria Balic (Institute of Oral Biology, University of Zurich) for critical reading and editing the manuscript.

## Authors contributions

Conceptualization: TAM, LV

Methodology: TAM, PP, HGR, AT, AM, FR

Investigation: PP, HGR, LLR, AT, TAM

Visualization: PP, HGR, AT, LV, TAM

Funding acquisition: TAM, AM

Project administration: TAM

Supervision: TAM, LV

Writing – original draft: HGR, LLR, PP, AT, AM, TAM

Writing – review & editing: HGR, LV, AT, AM, FR, TAM

## Data and materials availability

The accession number for all sequencing data reported in this paper is GEO: GSE198920; the reviewers can access the data using the token: mdmzaeqcbholbuf. All data will be made publicly available upon publication of the manuscript.

## Supplementary Materials

Supplementary material containing Materials and Methods, Supplementary Figures, Tables and References is provided in a separate document.

## Supplementary Materials

### Materials and Methods

#### Animals

Animal housing and experimentation were performed according to the Swiss Animal Welfare Law and in compliance with the regulations of the Cantonal Veterinary Office, Zurich (Licenses: 151/2014, 146/2017, 197/17). The animal facility provided standardized housing conditions, with a mean room temperature of 21 ± 1 °C, relative humidity of 50% ± 5%, and 15 complete changes of filtered air per hour (HEPA H 14filter); air pressure was controlled at 50 Pa. The light/dark cycle in the animal rooms was set to a 12 h/12 h cycle (lights on at 07:00, lights off at 19:00) with artificial light of approximately 40 Lux in the cage. The animals had unrestricted access to sterilized drinking water, and ad libitum access to a pelleted and extruded mouse diet in the food hopper (Kliba No. 3436; Provimi Kliba / Granovit AG, Kaiseraugst, Switzerland). Mice were housed in a barrier-protected specific pathogen-free unit and were kept in groups of max. 5 adult mice per cage in standard IVC cages (Allentown Mouse 500; 194 mm × 181 mm × 398 mm, floor area 500 cm2; Allentown, New Jersey, USA) with autoclaved dust-free poplar bedding (JRS GmbH + Co KG, Rosenberg, Germany). A standard cardboard house (Ketchum Manufacturing, Brockville, Canada) served as a shelter, and tissue papers were provided as nesting material. Additionally, crinklets (SAFE® crinklets natural, JRS GmbH + Co KG, Rosenberg, Germany) were provided as enrichment and further nesting material. The specific pathogen-free status of the animals was monitored frequently and confirmed according to FELASA guidelines by a sentinel program. The mice were free of all viral, bacterial, and parasitic pathogens listed in FELASA recommendations (*1*).

*K14Cre;Jagged1*^*fl/fl*^ conditional knockout mice were generated by crossing *K14:cre* (MGI: 2445832, Tg(KRT14-cre)1Amc/J(#004782)) and *Jagged1*^*flox*^ (MGI: 3577993, 129/Sv-Jag1<tm1Frad>) (*2*) mice. The animals were genotyped using the following primers: Cre Fw, 5’-CTG TTT TGC CGG GTC AGA AA-3’; Cre Rv, 5’-CCG GTA TTG AAA CTC CAG CG-3’; Jag1 Fw, 5’-GCA AGT CTG TCT GCT TTC ATC-3’; Jag1 Rv, 5’-AGG TTG GCC ACC TCT AAA TC-3’. The age of embryos was determined according to the vaginal plug (E0.5) and confirmed by morphological criteria. Animals were killed by cervical dislocation and E18.5 embryos were surgically removed and fixed overnight in 4% paraformaldehyde (PFA) in phosphate-buffered saline (PBS), pH 7.4. Newborn and adult animals were sacrificed by intracardiac perfusion with 4% PFA. Heads were then post-fixed in 4% PFA overnight at 4°C, thoroughly rinsed with PBS, and placed in 70% ethanol.

Embryos and newborn animals were washed in PBS, incubated in sucrose 30%, embedded in Tissue Tek® O.C.T.™ (4583, Sakura, Alphen aan den Rijn, Netherlands), and serially sectioned at 10 µm.

#### Dental nomenclature, imaging, measurements and phenotyping

The dental nomenclature used here is specific to murid rodents (Fig.1). M^n^ refers to the n^th^ upper molar, M_n_ to the n^th^ lower molar and Mn to both the n^th^ upper and lower molars. Cusps are respectively symbolized by a c^n^ and c_n_ for each upper and lower molar. Main cusps are numbered from 1 to 9 in upper molars and from 1 to 7 in lower molars from the mesio-lingual edge to the disto-vestibular edge of the tooth. High quality images of one wild-type mouse skull (D1404) and of two mutant mice skulls (D1437 and M12) were obtained using X-ray synchrotron microtomography at the European Synchrotron Radiation Facility (ESRF, Grenoble, France), beamline BM5, with a monochromatic beam at energy of 20 keV and using a cubic voxel of 7.45 µm. This method has been proven to be very useful for very precise imaging of small elements as teeth (*3*). 3D renderings were then performed using VG Studio Max 2.0 software. Dental morphological variations of location, size and interconnection of cusps were analyzed in the wild type and *K14Cre;Jagged1*^*fl/fl*^ mice (Supplemental material Fig. S2). Length (L) and width (W) for each molar were extracted at 0.001 mm using the LAS Core software (Leica^®^). The Student’s t-test and Fischer’s F-test was used on each dental measurement (W, L and d1-d5) for wild type and *K14Cre;Jagged1*^*fl/fl*^ mice to check mean and variance equality. In order to quantify the shape variations within M^1^ and M_1_ mesial parts, five other distances (d1-d5) were measured using LAS Core.

Overall tooth shapes were investigated by using an outline analysis. By registering the relative size and position of each cusp, this method appears suitable for tooth shape study. Fourier methods, notably Elliptic Fourier Transform (EFT), allow the description of complex outlines approximating them by a sum of trigonometric functions of decreasing wavelength (i.e. harmonics). x and y coordinates of 64 points equally spaced along dental outline were calculated to quantitatively describe the shape of M^1^ and M_1_. We applied EFTs to these data using EFA win software (*4*), extracting Fourier coefficients from the original outline, and normalizing these shape variables. This method considers the separate Fourier decomposition of the incremental change in x and y coordinates as a function of the cumulative length along the outline (*5*). For EFT, any harmonic n yields four Fourier coefficients: An and Bn for x, and Cn and Dn for y, which all contribute to describe the initial outline. We retained the first nine harmonics for M^1^ and the first five for M_1_, which represent the best compromise between measurement error and information content for these murine molars (*6*). However, the four coefficients of the first harmonic (A1–D1) were not included in the subsequent analyses because they were poorly discriminant and constituted background noise after the standardization step (size and orientation; (*6, 7*)).

#### Statistical analyses

We performed Student’s t-test and Fischer’s F-test on each dental measurement (W, L and D1-5) for WT and *K14Cre;Jagged1*^*fl/fl*^ mice to respectively check mean and variance equality. A principal component analysis (PCA) allowed evaluating a possible outline variation between WT and *K14Cre;Jagged1*^*fl/fl*^ mice. Variables were represented by the coefficients of each harmonics previously selected for M^1^ and M_1_ (respectively 32 and 16). A multivariate analysis of variations (MANOVA) allowed researching a potential significant difference between WT and *K14Cre;Jagged1*^*fl/fl*^ mice cohorts. This test included the coordinates of first axes of the PCA for which the sum met at least 95% of the total variation. These data were previously rank transformed since they did not fulfill the required parameters (i.e. normality, homoscedasticity of variances) for such statistical tests (*8*).

### RNA sequencing

#### Samples preparation

Lower molars were dissected from n = 4 E18.5 *K14cre; Jagged1*^*fl/fl*^ embryos and n = 4 *Jagged1*^*fl/fl*^ control littermates. Left and right lower first molars from the same animal were pooled. RNA was isolated using the RNeasy Plus Mini Kit (Qiagen AG, Switzerland) and subsequently purified by ethanol precipitation.

#### Library preparation

The quality of the isolated RNA was determined with a Qubit® (1.0) Fluorometer (Life Technologies, California, USA) and a Bioanalyzer 2100 (Agilent, Waldbronn, Germany). Only those samples with a 260 nm/280 nm ratio between 1.8–2.1 and a 28S/18S ratio within 1.5–2 were further processed. The TruSeq RNA Sample Prep Kit v2 (Illumina, Inc, California, USA) was used in the succeeding steps. Briefly, total RNA samples (100-1000 ng) were poly A enriched and then reverse-transcribed into double-stranded cDNA. The cDNA samples were fragmented, end-repaired and polyadenylated before ligation of TruSeq adapters containing the index for multiplexing Fragments containing TruSeq adapters on both ends were selectively enriched with PCR. The quality and quantity of the enriched libraries were validated using Qubit® (1.0) Fluorometer and the Caliper GX LabChip® GX (Caliper Life Sciences, Inc., USA). The product is a smear with an average fragment size of approximately 260 bp. The libraries were normalized to 10nM in Tris-Cl 10 mM, pH8.5 with 0.1% Tween 20.

#### Cluster Generation and Sequencing

The TruSeq PE Cluster Kit HS4000 or TruSeq SR Cluster Kit HS4000 (Illumina, Inc, California, USA) was used for cluster generation using 10 pM of pooled normalized libraries on the cBOT. Sequencing were performed on the Illumina HiSeq 4000 single end 125 bp using the TruSeq SBS Kit HS4000 (Illumina, Inc, California, USA).

#### Data analysis

Reads were quality-checked with FastQC. Sequencing adapters were removed with Trimmomatic (*9*) ^58^ and reads were hard-trimming by 5 bases at the 3’ end. Successively, reads at least 20 bases long, and with an overall average phred quality score greater than 10 were aligned to the reference genome and transcriptome of Mus Musculus (FASTA and GTF files, respectively, downloaded from Ensembl, GRCm38) with STAR v2.5.1 (*10*) ^59^ with default settings for single end reads.

Distribution of the reads across genomic isoform expression was quantified using the R package GenomicRanges (*11*) from Bioconductor Version 3.0. Differentially expressed genes were identified using the R package edgeR (*12*) from Bioconductor Version 3.0. A gene is marked as DE if it possesses the following characteristics: i) at least 10 counts in at least half of the samples in one group; ii) p <= 0.05; iii) fold change >= 0.5. Finally, gene sets were used to interrogate the GO Biological Processes database for an exploratory functional analysis. Contingency tables were constructed based on the number of significant and non-significant genes in the categories and we reported statistical significance using Fisher’s exact test.

#### Adapter Sequences

Oligonucleotide sequences for TruSeq™ RNA and DNA Sample Prep Kits

TruSeq Universal Adapter

5’

AATGATACGGCGACCACCGAGATCTACACTCTTTCCCTACACGACGCTCTTCCGATC T

TruSeq™ Adapters

TruSeq Adapter, Index 1

5’

GATCGGAAGAGCACACGTCTGAACTCCAGTCACATCACGATCTCGTATGCCGTCTTC TGCTTG

TruSeq Adapter, Index 2

5’

GATCGGAAGAGCACACGTCTGAACTCCAGTCACCGATGTATCTCGTATGCCGTCTTC TGCTTG

TruSeq Adapter, Index 3

5’

GATCGGAAGAGCACACGTCTGAACTCCAGTCACTTAGGCATCTCGTATGCCGTCTTC TGCTTG

TruSeq Adapter, Index 4

5’

GATCGGAAGAGCACACGTCTGAACTCCAGTCACTGACCAATCTCGTATGCCGTCTTC TGCTTG

TruSeq Adapter, Index 5

5’

GATCGGAAGAGCACACGTCTGAACTCCAGTCACACAGTGATCTCGTATGCCGTCTTC TGCTTG

TruSeq Adapter, Index 6

5’

GATCGGAAGAGCACACGTCTGAACTCCAGTCACGCCAATATCTCGTATGCCGTCTTC TGCTTG

TruSeq Adapter, Index 7

5’

GATCGGAAGAGCACACGTCTGAACTCCAGTCACCAGATCATCTCGTATGCCGTCTTC TGCTTG

TruSeq Adapter, Index 8

5’

GATCGGAAGAGCACACGTCTGAACTCCAGTCACACTTGAATCTCGTATGCCGTCTTC TGCTTG

TruSeq Adapter, Index 9

5’

GATCGGAAGAGCACACGTCTGAACTCCAGTCACGATCAGATCTCGTATGCCGTCTTC TGCTTG

TruSeq Adapter, Index 10

5’

GATCGGAAGAGCACACGTCTGAACTCCAGTCACTAGCTTATCTCGTATGCCGTCTTC TGCTTG

TruSeq Adapter, Index 11

5’

GATCGGAAGAGCACACGTCTGAACTCCAGTCACGGCTACATCTCGTATGCCGTCTTC TGCTTG

TruSeq Adapter, Index 12

5’

GATCGGAAGAGCACACGTCTGAACTCCAGTCACCTTGTAATCTCGTATGCCGTCTTC TGCTTG

TruSeq Adapter, Index 13 5’GATCGGAAGAGCACACGTCTGAACTCCAGTCACAGTCAACAATCTCGTATGCCGT CTTCTGCTTG

TruSeq Adapter, Index 14

5’

GATCGGAAGAGCACACGTCTGAACTCCAGTCACAGTTCCGTATCTCGTATGCCGTCT TCTGCTTG

TruSeq Adapter, Index 15

5’

GATCGGAAGAGCACACGTCTGAACTCCAGTCACATGTCAGAATCTCGTATGCCGTCT TCTGCTTG

TruSeq Adapter, Index 16

5’

GATCGGAAGAGCACACGTCTGAACTCCAGTCACCCGTCCCGATCTCGTATGCCGTCT TCTGCTTG

TruSeq Adapter, Index 18 4

5’

GATCGGAAGAGCACACGTCTGAACTCCAGTCACGTCCGCACATCTCGTATGCCGTCT TCTGCTTG

TruSeq Adapter, Index 19

5’

GATCGGAAGAGCACACGTCTGAACTCCAGTCACGTGAAACGATCTCGTATGCCGTCT TCTGCTTG

TruSeq Adapter, Index 20

5’

GATCGGAAGAGCACACGTCTGAACTCCAGTCACGTGGCCTTATCTCGTATGCCGTCT TCTGCTTG

TruSeq Adapter, Index 21

5’

GATCGGAAGAGCACACGTCTGAACTCCAGTCACGTTTCGGAATCTCGTATGCCGTCT TCTGCTTG

TruSeq Adapter, Index 22

5’

GATCGGAAGAGCACACGTCTGAACTCCAGTCACCGTACGTAATCTCGTATGCCGTCT TCTGCTTG

TruSeq Adapter, Index 23

5’

GATCGGAAGAGCACACGTCTGAACTCCAGTCACGAGTGGATATCTCGTATGCCGTCT TCTGCTTG

TruSeq Adapter, Index 25

5’

GATCGGAAGAGCACACGTCTGAACTCCAGTCACACTGATATATCTCGTATGCCGTCT TCTGCTTG

TruSeq Adapter, Index 27

5’

GATCGGAAGAGCACACGTCTGAACTCCAGTCACATTCCTTTATCTCGTATGCCGTCT TCTGCTTG

### RNA isolation real time-PCR

For real time PCR analysis, lower molars were dissected from n= 8 E18.5 *K14cre; Jagged1*^*fl/fl*^ embryos and n = 8 *Jagged1*^*fl/fl*^ control littermates. Left and right lower first molars from the same animal were pooled. RNA was isolated using the RNeasy Plus Mini Kit (Qiagen AG, Switzerland) and subsequently purified by ethanol precipitation. Reverse transcription of the isolated RNA was performed using the iScript™ cDNA synthesis Kit and according to the instructions given (Bio-Rad Laboratories AG, Cressier FR, Switzerland). Briefly, 1000 ng of RNA were used for reverse transcription into cDNA. Nuclease-free water was added to add up to a total of 15 μl. 4 μl of 5x iScript reaction mix and 1 μl of iScript reverse transcriptase were added per sample in order to obtain a total volume of 20 μl. The reaction mix was then incubated for 5 min at 25°C, for 30 min at 42°C and for 5 min at 85°C using a Biometra TPersonal Thermocycler (Biometra AG, Göttingen, Germany). The 3-step quantitative real-time PCRs were performed using an Eco Real-Time PCR System (Illumina Inc., San Diego CA, USA).

The reaction mix was composed of 5 μl of SYBR® Green PCR Master Mix reverse and forward primers (200 nM), and 2 ng of template cDNA. The thermocycling conditions were: 95°C for 10 min, followed by 40 cycles of 95°C for 15 sec, 55°C for 30 sec and 60°C for 1 min. Melt curve analysis was performed at 95°C for 15 sec, 55°C for 15 sec and 95°C for 15 sec. Expression levels were calculated by the comparative ΔCt method (2−ΔCt formula), normalizing to the Ct-value of the *36B4* housekeeping gene.

### Immunohistochemistry

Cryosections were air dried for 1 hour at room temperature, then washed with PBS to remove excess of Tissue Tek® O.C.T.™. Endogenous peroxidases were inhibited by incubating the sections in a solution composed of 3% H_2_O_2_ in Methanol at −20°C for 20 minutes. Specimens were then blocked with PBS supplemented with 2% fetal bovine serum and thereafter incubated with primary antibodies for 1 hour at room temperature. The following primary antibodies were used: Rabbit anti-Notch1-ICD (1:50, ab8925, Abcam, Cambridge, United Kingdom), Rabbit anti-Hes1 (1:50, 19988, Cell Signaling, Danvers, MA, USA), Rabbit anti-Hes5 (1:50, ab194111, Abcam, Cambridge, United Kingdom), Rabbit anti-β-catenin (1:50, 8480S, Cell Signaling, Danvers, MA, USA), Rabbit anti-Ki67 (1:100, ab15580, Abcam, Cambridge, United Kingdom), Rabbit anti-Amelogenin (1:100, ab153915, Abcam, Cambridge, United Kingdom). For negative controls, primary antibody was omitted. The sections were then incubated with a biotinylated secondary antibody (Vector Vectastain ABC kit PK-4001-1; Vector Laboratories LTD, Peterborough, UK). Sections were then incubated with AEC (3-amino-9-ethylcarbazole; AEC HRP substrate Kit - SK4200; Vector Laboratories LTD, Peterborough, UK) to reveal the staining, counterstained with Toluidine Blue, mounted with Glycergel (C0563, Agilent Technologies, Santa Clara, CA, USA) and imaged with a Leica DM6000 light microscope (Leica Microsystems, Schweiz AG, Heerbrugg, Switzerland).

### 3D morphing of tooth geometries and computation/prediction of human Jagged1-mutant M^1^

The 3D morphing of all tooth geometries was conducted in ANSA by BETA CAE Systems A.G. (CH-6039 Switzerland) and the methodological approach can be broken down into the following 5 steps.

#### Tooth surface generation

Average tooth morphologies were computed for the M^1^ of: WT rodent (based on two characteristic teeth), a mutant one (considering nine Jagged1-mutant crown morphologies) and a human M^1^ (resulting from 246 molars). This process facilitated the detection and association of the prevalent features, to compute realistic reference models. Especially for the human molar M^1^ a large existing data base was available, which was used as the starting point of calculating the respective average tooth morphology. The dataset is based on a set of existing impressions of carious-free and intact tooth surfaces of West European young people within the age of 9 to 20 years. From the impressions, stone replicas were made, and the clinical visible parts of their surfaces were measured with a 3D-scanning device. The resolution of the measuring process was 50 µm x 50 µm (x,y). All data sets were aligned in position and orientation within the same coordinate system: For this, a representative molar with an appropriate orientation was chosen ^62^. All other molars were superimposed with this representative molar by a least-square fitting routine (Match3D) (*13-15*). This guaranteed that all occlusal features had the same orientation. The pattern of cusps and grooves varies considerably across individuals, even though some morphological properties, such as the overall layout of the main cusps and fissures, are shared by all samples. These common features allow to establish correspondence between the scans z(x,y) with a modified optical flow algorithm in an automated procedure (*16*). Based on this correspondence, the *x, y, z* - coordinates can be averaged and result in a typical representation of the human M^1^, which was used further in this process (*13, 14*).

#### Pre-processing

Prior to applying the morphing algorithms to the teeth, proxy geometries were created, by scaling all teeth (mouse WT and mutant M^1^ and human M^1^) to comparable dimensions. Teeth were then aligned based on characteristic morphological patterns (e.g. tooth cusps and grooves). Mouse and human teeth were positioned by a point-to-point spatial relation of their functional features: the 4 cusps of the human M^1^ (protocone, metacone, paracone, hypocone) were aligned to the 4 distal cusps (c^5^, c^6^, c^8^, c^9^) of the mouse M^1^.

#### Creation of a dense 3D correspondence (Features mapping)

The non-uniform triangulated meshes of the *.stl files, were used as bi-linear maps for the morphing algorithm, facilitating the determination of a dense correspondence across both source (WT) and target (mutant) tooth geometry. The mouse mutant M^1^ consisted of 249.797 triangular elements whereas the mouse WT M^1^ and human M^1^, of 252.296 and 51.476 respectively. A correspondence was algorithmically established, by identifying the minimum projected distance between each node of the source mesh to a node or an interpolation of multiple nodes on the target geometry. Source nodes that are not paired with correspondences, are handled as “in-between positions” and placed as simple linear interpolations of the vertex positions in the target mesh. The use of these nodes in the mesh triangulation, increases the quality of the morph function, despite not being visible on the target.

To provide a reference plane for the translation matrix, pairs of points were placed on the source and target geometry to identify locations that should be in correspondence. These boundary locations were selected below the functional surfaces of the molars (in proximity to the cervical margin line) to ensure the unobstructed morphing of the molar crown morphology. Areas of interest were isolated based on the statistically significant variations found in mouse WT M^1^ vs mutant M^1^. The computed transformation matrix was then applied to the 3D data set of the human M^1^, to predict the crown morphology in individuals carrying Jagged1 mutations.

#### Construction of the 3D morphs

Polygonal surface approaches, such as 3D morphing, require mapping of the characteristic topological landmarks (as described above), followed by re-meshing of the morphed surfaces to achieve a realistic convergence. The quality-oriented reconstruction of the source model’s grid produces a robust 3D morph.

#### Application of the 3D morphs to human molars

The transformation vectors required to morph the human healthy M^1^ into a M^1^ in individuals carrying Jagged1 mutations were formulated as an algorithmic matrix. This matrix was then stored as a geometry-independent parameter, reflecting the differences of the crown surface between WT and mutant M^1^. Transferring the deformation map of this parameter to the 3D data set of the human M^1^ (average model) facilitates the computation of the M^1^ crown morphology, which is expected to be representative both in terms of morphology and size of the human mutant M^1^.

## Supplementary Figures

**Supplementary Figure 1.**
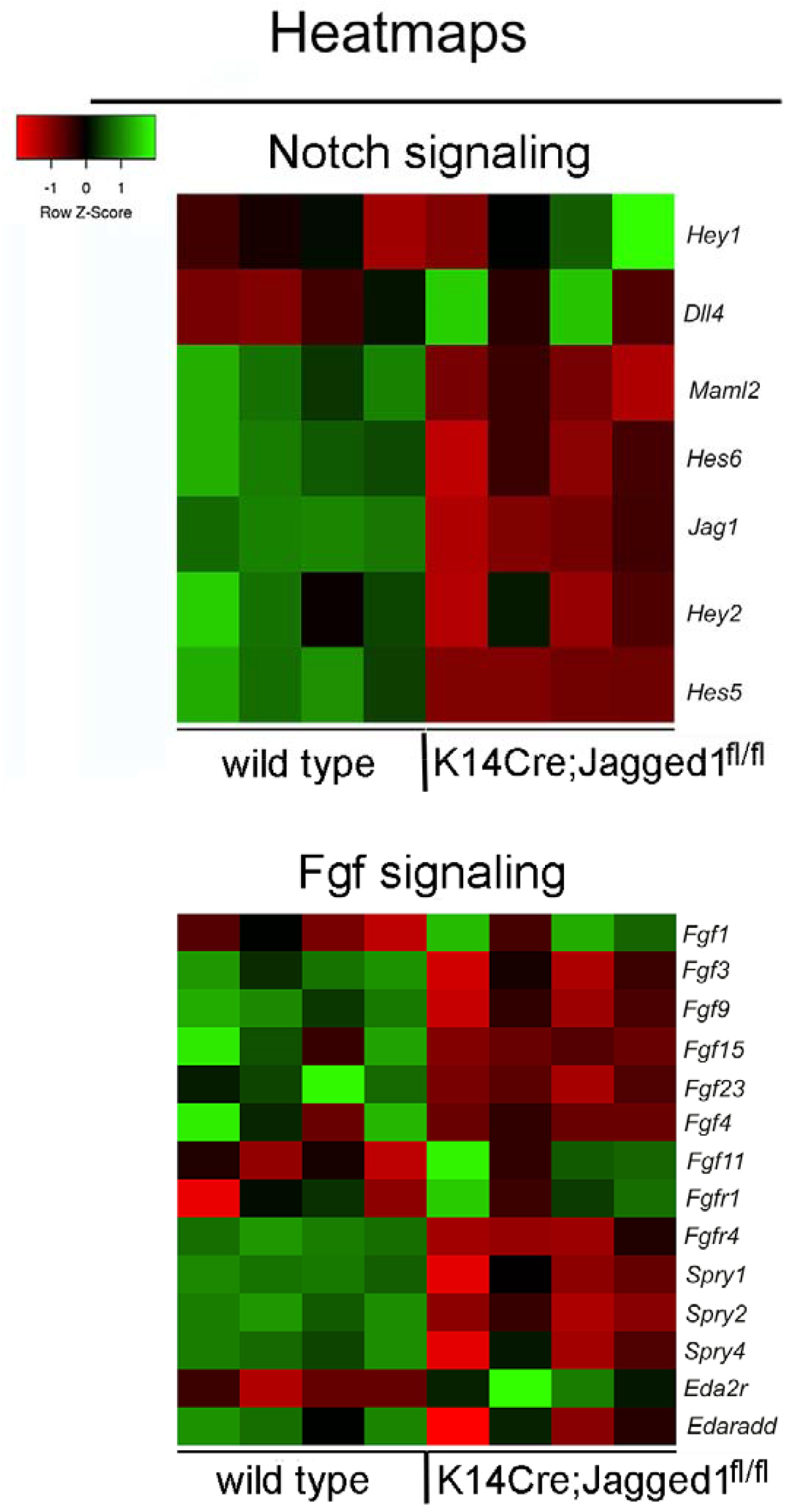
RNA sequencing analysis of first molars isolated from E18.5 WT and *K14Cre;Jagged1*^*fl/fl*^ mouse embryos. Heatmaps showing the main deregulated genes involved in the Notch and FGF signaling pathways in *K14Cre;Jagged1*^*fl/fl*^ molars.

